# Validating layer-specific VASO across species

**DOI:** 10.1101/2020.07.24.219378

**Authors:** Laurentius Renzo Huber, Benedikt A Poser, Amanda L Kaas, Elizabeth J Fear, Sebastian Desbach, Jason Berwick, Rainer Goebel, Robert Turner, Aneurin J Kennerley

**Affiliations:** MBIC, Department of Cognitive Neuroscience, Faculty of Psychology and Neuroscience, Maastricht University, The Netherlands; Department of Chemistry, University of York, York, United Kingdom; Department of Psychology, University of Sheffield, Sheffield, United Kingdom; Neurophysics Department Max Planck Institute for Human Cognitive and Brain Sciences, Leipzig, Germany and Sir Peter Mansfield Imaging Centre, University of Nottingham, Nottingham, United Kingdom

**Author notes:** contributed equally (Laurentius Renzo Huber, Aneurin J Kennerley).

**Keywords:** fMRI, laminar, layer, pre-clinical, sub-millimeter, somatosensory stimulation, draining vein, depth-dependent fMRI, sub-millimeter, concurrent imaging, Optical Imaging Spectroscopy, cerebral blood volume, MION, VASO

## Abstract

Cerebral blood volume (CBV) has been shown to be a robust and important physiological parameter for quantitative interpretation of functional (f)MRI, capable of delivering highly localized mapping of neural activity. Indeed, with recent advances in ultra-high-field (>=7T) MRI hardware and associated sequence libraries, it has become possible to capture non-invasive CBV weighted fMRI signals across cortical layers. One of the most widely used approaches to achieve this (in humans) is through vascular-space-occupancy (VASO) fMRI. Unfortunately, the exact contrast mechanisms of layer-dependent VASO fMRI have not been validated and thus interpretation of such data is confounded. Here we cross-validate layer-dependent VASO fMRI contrast in a preclinical rat model using well established (but invasive) imaging methods in response to neuronal activation (somatosensory cortex) and respiratory challenge (hypercapnia). In particular VASO derived CBV measures are directly compared to concurrent measures of total haemoglobin changes from high resolution intrinsic optical imaging spectroscopy (OIS). Through direct comparison of response magnitude, across time, negligible changes in hematocrit ratio during activation (neuronal or vascular) are inferred. Quantified cortical layer profiling is demonstrated and in agreement between both VASO and contrast enhanced fMRI (using monocrystalline iron oxide nanoparticles, MION). Responses show high spatial localisation to layers of cortical excitatory and inhibitory processing independent of confounding large draining veins which hamper BOLD fMRI studies. While we find increased VASO based CBV reactivity (3.1 ± 1.2 fold increase) in humans compared to rats it is demonstrated that this reflects differences in stimulus design rather than confounds of the VASO signal source. Together, our findings confirm that the VASO contrast is indeed a reliable estimate of layer-specific CBV changes. This validation study increases the neuronal interpretability of human layer-dependent fMRI results and should supersede BOLD fMRI as the method of choice in neuroscience application studies.

**Highlights:**

- Our goal is to validate layer-specific VASO fMRI with gold standard methods
- Layer-specific VASO sequences are implemented for 7T imaging in humans and rats
- Comparisons of VASO, optical imaging, and MION confirm the expected contrast origin
- Somatosensory stimulation in humans and rats reveal the same layer-fMRI signatures
- We confirm that VASO is a valid measure to estimate layer-specific neural activity

**Graphical abstract:** 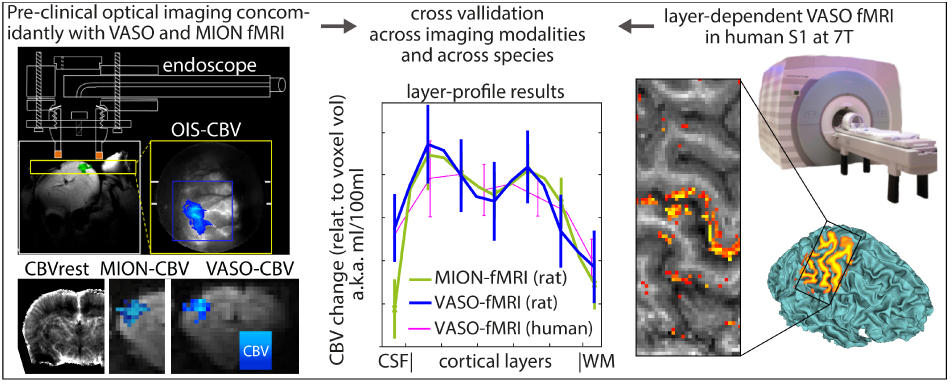

## 1. Introduction

The layered architecture of the cerebral cortex was first identified in the mid-late 1800s. The hierarchical structure of the cortex has since been parcellated based upon cell type, and the density of either cell bodies (Brodmann 1909; Economo and Koskinas 1925) or myelinated fibres (Smith 1907; Vogt and Vogt 1919). It is noted that the layering schemes based upon cyto- or myeloarchitecture methodologies do differ (Lashley and Clark 1946); and modern delineation of layer boundaries now follow a multiparametric approach (Schleicher et al. 2005). While the definition of laminar borders moves towards statistical inference, it is widely accepted that the laminar organisation of cells and associated afferents is central to cortical function (Douglas and Martin 2004a); with feed-forward and feedback pathways having known layer-specific termination patterns (Douglas and Martin 2004b; Larkum et al. 2018). Non-invasive imaging of layer dependent functional activity supplements our understanding of structural parcellation and will result in a rapid increase of our knowledge of the working human brain. Technological advances in MRI, both in terms of magnetic field (Polimeni and Uludağ 2018), imaging gradient coil strength, detection hardware and advanced sequence designs (Poser and Setsompop 2018), have permitted high resolution, in-vivo imaging of cortical anatomy (Trampel et al. 2019; Turner 2013). Conventional MR measurements do indeed reflect the cortical architecture found in earlier histological studies (Eickhoff et al. 2005). The recent marriage of such data with methodological advances in quantitative MRI (based on phase contrast (Duyn et al. 2007) and diffusion (Assaf 2019; Leuze et al. 2014) weighted imaging) delivers robust mapping of cortical layer structure.

However, complementary mapping of brain function across the cortical layers with Blood Oxygenation Level Dependent (BOLD) functional (f)MRI, falls behind. A major reason for this is that conventional 2D gradient-echo (GE) BOLD measurements are confounded by signals originating from the large draining veins, driven by both large magnetic susceptibility effects and magnetic field angle dependence (Chu et al. 1990; Fracasso et al. 2018; Gagnon et al. 2015). Thus functional maps, while locally specific, show signals spreading across cortical layers and columns (Kennerley et al. 2005; Turner 2002). Equally, conventional 2D single-shot EPI, Spin-echo (SE) BOLD techniques, while thought to be more specific to the microvasculature, and so better suited for the study of cortical columns (Koopmans and Yacoub 2019; Norris 2012), still demonstrate some weighting to the pial vessels (Goense and Logothetis 2006) due to significant intravascular dephasing effects (Boxermann et al. 1995; Martindale et al. 2008).

To overcome these limitations and approach the mesoscopic spatial regime of cortical layers for brain function, alternative non-BOLD quantitative contrast mechanisms are required (Chai et al. 2019; Huber et al. 2019) in combination with modern hardware and readout regimes (Norris and Polimeni 2019; Yacoub and Wald 2018) - akin to the advances in structural based imaging.While an extensive range of techniques exists (e.g. GRASE (Feinberg et al. 2008; Oshio and Feinberg 1991), ASL (Ivanov et al. 2016), SSFP (Goa et al. 2014; Scheffler et al. 2018), functional diffusion (Truong and Song 2009), phase-sensitive fMRI (Menon 2002; Vu and Gallant 2015) etc); one of the fastest growing approaches for layer-based fMRI utilises slice-saturation slab-inversion vascular space occupancy (SS-SI VASO)(Huber et al. 2014b).The VASO contrast exploits the difference between longitudinal relaxation times (T_1_) of tissue and blood (Lu et al. 2003). Contrast is generated by applying a magnetization inversion pulse before signal acquisition, to effectively null the contribution of blood water pool at the time of signal excitation, while maintaining substantial tissue signal for detection. Thus decreases in MR signal intensity during neuronal activity reflect the associated hemodynamic increase in the volume fraction of blood (or cerebral blood volume, CBV) (Attwell and Iadecola 2002) within the imaged voxel.

VASO based CBV measurements are particularly attractive for layer-specific functional brain imaging in humans because theoretically the method:

I. offers a quantitative measure, being described in meaningful physical units (ml);
II. it is believed to be insensitive to large draining veins (Lu et al. 2013);
III. promises a more robust pseudo-measure of neuronal activity, maintaining signal change even in light of significant physiological changes (which have been demonstrated to dramatically affect BOLD fMRI (Kennerley et al. 2012a)).

Slice selective slab-inversion (SS-SI) VASO is widely used across the literature for non-invasive imaging of microvascular layer-specific signal responses (Beckett et al. 2020; Chai et al. 2019; Finn et al. 2019; Guidi et al. 2017; Huber et al. 2018; Kurban et al. 2020; Persichetti et al. 2020; Yang and Yu 2019; Yu et al. 2019).

Although SS-SI-VASO is recognised as a powerful tool for layer based fMRI studies, the ‘exact’ signal mechanisms remain enigmatic. Indeed over the last decade, much focus and debate has centred on enhancing our understanding of the VASO signal source (Donahue et al. 2006; Hua et al. 2009; Jin and Kim 2008; Scouten and Constable 2007; 2008; Uh et al. 2011; Wu et al. 2010). While the inverse relationship between VASO contrast and cerebral blood volume (CBV) changes has been partially validated, with Positron Emission Tomography (Uh et al. 2011) and contrast agent enhanced MRI (Jin and Kim 2006; 2008; Lin et al. 2011; Lu et al. 2005), it is important to note that some of these comparison studies were of low spatial resolution (e.g. 3 x 3 x 3 mm3), used excessive smoothing filters, ignored some of the more interesting temporal dynamics of the response, and were based on non-quantitative correlative analysis. VASO contrast could be confounded by the complex interplay between cerebral blood volume, flow and extravascular BOLD effects (with inflow of fresh spins and partial volume effects further confounding measurement) (Donahue et al. 2006). Thus the high correlations between VASO and alternative imaging methods may be erroneously dominated by these non-CBV components. Also, the layer-specific application of the VASO contrast remains unvalidated. The missing validation with gold standard methods might limit the wider uptake of the SS-SI VASO method for high-resolution fMRI.

To investigate whether the layer-dependent VASO response is indeed directly driven by stimulus induced changes in CBV, there is an urgent need for improved validation studies with particular focus on signal quantification (in physical units of ml) and detailed study of the signal contribution from large veins. While the insensitivity of high resolution layer-based VASO contrast to large draining vein effects has been qualitatively compared with standard GE-BOLD (reviewed in (Huber et al. 2019)), it has not been quantitatively validated with independent ‘ground-truth’ CBV imaging modalities to date.

Here we cross-validate layer-dependent VASO fMRI contrast with well established (but invasive) high resolution imaging methods (both optical and Fe3+ contrast based CBV-weighted MRI) in a pre-clinical rat model in response to neuronal activation (somatosensory cortex) and respiratory challenge (hypercapnia). The leading hypothesis is that these independent methodologies will offer a deeper understanding/corroboration of the signal source of VASO contrast across the cortical layers; and permit in-depth study of the interspecies differences in VASO contrast. Optical imaging spectroscopy allows the assessment of haemoglobin concentration changes (in the two oxygenation states; summed to give total haemoglobin); while CBV-weighted MRI (comparable with VASO) actually assesses plasma volume. This study uses our specialised concurrent imaging based approach (Kennerley et al. 2012b) for improved and direct comparison between methodologies. Inferences about hematocrit changes (Hall et al. 2014) during neuronal activation and the influence on the measured signal magnitude (Levin et al. 2001) can therefore be made.

As alluded to above, when considering CBV fMRI contrast there are known interspecies differences which raise debate across the fMRI research field. There are two specific features of pre-clinical CBV-weighted fMRI that have not been confirmed in equivalent human studies to date. These discrepancies concern i) the absolute magnitude of CBV changes and ii) the overall temporal dynamics. Human CBV sensitive VASO fMRI studies find blood volume changes in the order of 30 - 90% (compared to CBV_*r*_*est*) in response to both visual and sensorimotor based neuronal activation (Donahue et al. 2006; Gu et al. 2006; Hua et al. 2009; Huber et al. 2014b; Lu et al. 2003; 2013; 2004b;c; Poser and Norris 2007; Scouten and Constable 2008; Shen et al. 2009). Comparable animal studies with invasive methodologies report much smaller CBV changes of 10% - 20% (Mandeville et al. 1998), *≈* 5-8% (Kennerley et al. 2005), 5% (Kennerley et al. 2012a), and 4% - 7% (Lu et al. 2004b) across a broad range of somatosensory stimuli and associated durations. Independent of such divergences in absolute CBV change, time course dynamics are also often qualitatively different between human VASO studies and equivalent measures reported across the animal literature. Human VASO studies suggest a fast recurrence to baseline of CBV after stimulus cessation (in the range of 10 - 15 s) (Hua et al. 2011; Lu et al. 2004a; Poser and Norris 2007; van Zijl et al. 2012), while rat data demonstrates delayed compliance (with a post-stimulus signal return to baseline in the order of 30 - 60 s) (Kennerley et al. 2005; Kida et al. 2007; Kong et al. 2004; Mandeville et al. 1999).

It can be argued that the interspecies differences in CBV amplitude and temporal dynamics (outlined above) reflect i) obvious methodical differences and thus signal source confounds, ii) task related and natural response differences, or simply iii) unavoidable anesthesia-dependent confounds in pre-clinical models. Extensive studies have suggested that anesthesia can ‘slightly’ reduce the overall response amplitude, but use of these agents cannot explain the temporal CBV dynamic differences between animals and humans (Berwick et al. 2002; Martin et al. 2006; Sicard et al. 2003; Zong et al. 2012). An extensive interspecies VASO contrast comparison between humans and animals, underpinned with data from more invasive imaging methods in the latter model, will help resolve these differences and aid our interpretation of the VASO fMRI signal source. VASO fMRI is a non-invasive contrast, which can easily be applied both in rats and in humans. Here we will investigate the haemodynamic response to stimulation of the primary somatosensory region of both the human and rat cortex (subsequently avoiding potential brain region confounds). In addition we will investigate CBV changes in response to hypercapnic respiratory challenges. It is hypothesised that similar hemodynamic control pathways in humans and rats exist for such challenges (Mortola and Lanthier 1996), enabling quantitative comparisons (after controlling for body mass and baseline blood oxygen saturation (Gray and Steadman 1964)). Despite initial VASO comparisons with CBV-weighted fMRI at high spatial resolution in monkey visual cortex (Goense et al. 2012; Huber et al. 2014a) and cat visual cortex (Jin and Kim 2006; 2008), the above described difference of response magnitudes between VASO and non-VASO CBV imaging have not been resolved to date.

The present study therefore aims to investigate the cortical layer based sensitivity of VASO-derived CBV measurements in both human and rat models. VASO insensitivity to changes in the large draining veins on the cortical surface will be confirmed in rodents by direct comparison of functional changes (to neuronal activation/hypercapnia) acquired by VASO, contrast enhanced/CBV weighted fMRI and BOLD fMRI, with concomitantly acquired optical measures of total haemoglobin and blood oxygenation. Data will be used to validate the layer-dependent CBV profile of VASO and contrast enhanced MRI within the same animals, within the same session. Ultimately, we seek to compare layer-dependent fMRI profiles of CBV and BOLD across rodents and humans using the same non-invasive SS-SI-VASO sequence in primary somatosensory cortex measured at the same MRI field strength (7 T). This high-resolution validation study of VASO contrast with an established preclinical model will be of particular interest to the growing field of high-resolution layer based fMRI; data will deepen understanding of the VASO fMRI signal source and help increase the wider uptake of the SS-SI-VASO method.

Here, we present an fMRI analysis software suite LAYNII that is specifically designed for layer-fMRI data and that addresses all of the above listed challenges. LAYNII is designed to perform layer analyses entirely in voxel space and consists of many modular programs that perform layer-fMRI specific tasks:

## 2. Methods

Here we implement VASO based fMRI in a preclinical model utilising concurrent intrinsic optical imaging techniques (**?**) (developed over 15 years ago) to investigate the underlying haemodynamics with high spatio-temporal resolution.

VASO, as a potential measure of cerebral blood volume changes in response to neuronal activation, was originally developed for use in the human brain at low magnetic fields (1.5 T) (Lu and van Zijl 2012). Unfortunately, as magnetic field strength increases the T_1_ relaxation time constants for intravascular and extravascular compartments converge (Jin and Kim 2008); and thus contrast to noise between nulled and non-nulled signals decreases. To account for this convergence and make the VASO sequence viable (in terms of contrast to noise) at higher field strengths (7T+), a variant SS-SI VASO pulse sequence was previously introduced (Huber et al. 2014b). For application in an animal model at high fields (7T+), distinct features in physiology (e.g. short arterial arrival time (Calamante et al. 1999)) and associated magnetization dynamics must be reconsidered. Due to the fast heart rate in rat models there is increased risk of inflow of non-inverted blood magnetization into the imaging slice during the blood nulling time, and thus if unaccounted for, VASO signal can become unreliable (Hua et al. 2013) as steady state magnetisation is impaired (Donahue et al. 2009). To correct for this difference, additional modules to control the magnetization were introduced to the SS-SI-VASO sequence here. These include implementation of additional slice-selective and global spin-reset saturation pulses to account for the different z-magnetization relaxation histories (steady-state) of stationary tissue and blood flowing into the imaging slice (Hua et al. 2013; Lu 2008). This is used to increase functional contrast to noise and avoid contamination of changes in CBF. One repetition time (TR) of the adapted SS-SI-VASO sequence is depicted in Fig. 1A.

**Figure 1.**
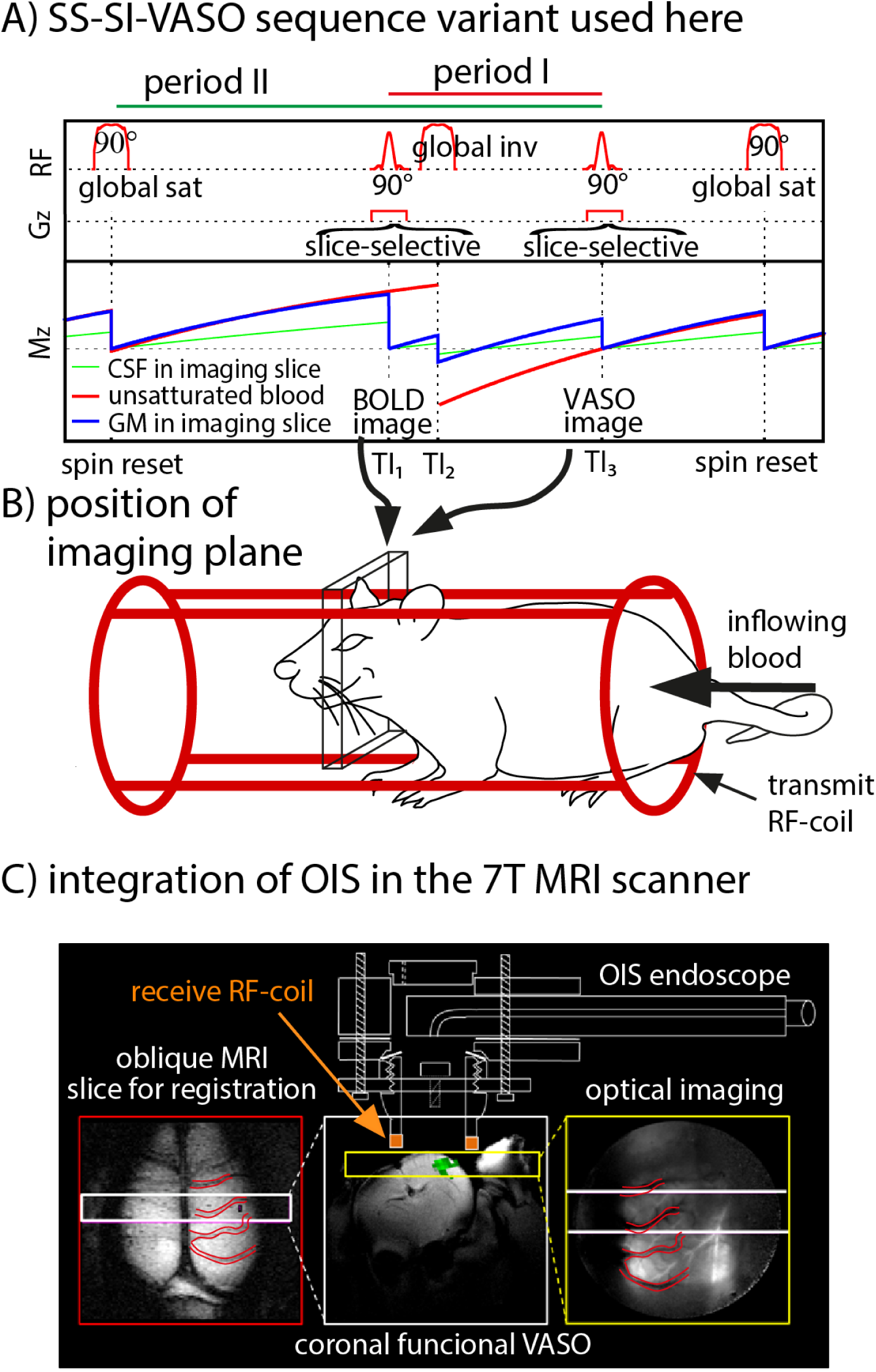
Acquisition procedure of SS-SI VASO used in rats to account for the faster arterial arrival time compared to humans. Acquisition procedure of SS-SI VASO in rats: A) depicts a pulse sequence diagram of SS-SI VASO modified for application in rodent brains at high fields (7T+). Before every TR, a global adiabatic saturation pulse is applied, ensuring control of the steady state magnetization steady-state across the parts of the animal covered by the large quadrature volume resonator. This global reset pulse shortens the time that the flowing blood pool magnetization requires to transition into a steady-state. In-plane magnetization is saturated by a slice-selective 90*°*excitation pulse prior to the acquisition of a BOLD weighted image (TI_1_; without blood nulling). After this saturation pulse, a global inversion pulse is applied (TI_2_) and subsequent VASO signal acquisition performed at the blood-nulling time TI_3_. B) depicts the approximate position of slice-selective saturation and ‘global’ inversion pulses. A large 180mm inner diameter quadrature coil is used to ensure inversion across the body and enable concurrent optical imaging setup. C) depicts a schematic illustration of our concurrent fMR and intrinsic optical imaging spectroscopy (OIS) setup and acquisition planes. Concurrent optical imaging inside the bore of the scanner is permitted through a non-magnetic medical endoscope. The apparatus to hold the endoscope in place is surgically attached to the cranium and has a built-in surface coil for MR RF reception. The large draining veins are visible in both 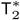 weighted MR images and corresponding high-resolution optical images. Such vascular landmarks help co-register data between the imaging modalities.

Before every TR, a global adiabatic 90*°* spin-reset pulse is applied to eliminate the spin history of all magnetization pools (e.g. stationary tissue, nulled/un-nulled blood etc.) within the 1H quadrature resonator (Bruker 1P-T9561, 300MHz, 1kW max, 200/180mm OD/ID) used for RF transmission (Fig. 1B). The amplitude and phase shape-functions of this radio-frequency pulse were adapted from the TR-FOCI pulse class, designed for use at 7 T (Hurley et al. 2010). In order to convert the original TR-FOCI pulse from an inversion pulse to a 90*°* saturation pulse, a 90*°* phase skip of the pulse amplitude was introduced half-way through the pulse duration, resulting in an inversion efficiency of 50%, despite the B_1_ inhomogeneities of the transmit coil (Mispelter et al. 2006). A more detailed description on the operating principle of this RF pulse can be found in (Huber et al. 2014b). All blood in the imaging slice is expected to be nulled to provide a pure VASO contrast, when two assumptions are fulfilled: firstly, the microvasculature of the imaging slice is refilled during period I (Fig. 1A)-blood velocity and transit time measurements suggest that the microvasculature of a 2 mm imaging slice is refilled in less than 1.5 s (Hutchinson et al. 2006; Ivanov et al. 1981; Kennerley et al. 2010; Kim and Bandettini 2010; Kleinfeld et al. 1998; Pawlik et al. 1981)–and secondly, no fresh (uninverted) blood from the hindquarters of the animal flows into the imaging slice during period II (Fig. 1A).

### 2.1. Experimental setup for data acquisition in rats

All aspects of these concurrent imaging methods and their development have been previously described (Kennerley et al. 2012b). A brief description is given here for completeness. All experiments were performed with UK Home Office approval under the Animals (Scientific Procedures) Act of 1986. Female hooded Lister rats (total n = 5) weighing (250–400 g) were kept in a 12 h dark/light cycle, at a constant temperature of 22 °C, with food and water ad libitum. Animals were anesthetized (i.p. urethane 1.25 g/kg) with additional 0.1 ml doses administered as necessary. A single subcutaneous injection of Atropine (0.4 mg/kg) reduced mucous secretions throughout the experiment. Rectal temperature was maintained at 37°C throughout surgical and experimental procedures using a homeothermic blanket (Harvard Apparatus Inc, USA). Animals were tracheotomized to allow artificial ventilation and permit controlled respiratory challenges. Ventilation parameters were adjusted to maintain blood gas measurements (measured via blood letting) within physiological limits (pO2 = 105 mm Hg ± 4; pCO2 = 38 mm Hg ± 5). Left and right femoral veins and arteries were cannulated. Phenylephrine (0.13–0.26 mg/h) was infused i.v. to maintain blood pressure (measured via pressure transducer connected to the arterial cannulae - CWE systems Inc. USA) between physiological limits (MABP, 100–110 mm Hg (Nakai and Maeda 1999)).

Intrinsic optical imaging in rat models is an invasive methodology requiring direct access to the cortex for spectroscopic analysis of remitted light (Lieke et al. 1989; Mayhew et al. 1999). To facilitate this, animals were placed in a stereotaxic frame (Kopf Instruments, USA). The skull overlying the right somatosensory cortex was thinned to translucency with a dental drill under constant cooling with saline. A bespoke single loop RF surface coil, integrated into a 20 mm diameter Perspex well was fixed to the animals’ head using dental cement, ensuring that the thin window lay in the centre of the well. This well is used to both secure the animal inside the MRI scanner and also holds the non-magnetic endoscope (Endoscan Ltd, London) used to permit concurrent optical imaging of the cortex within the magnet bore (Fig. 1C). The well is filled with deuterium oxide (D_2_O) to i) reduced optical specularities from the skull surface (for optical imaging); and ii) reduce air-tissue susceptibility artefacts (around the thinned cranial window for high field fMRI) whilst being MR silent at the proton resonant frequency.

Platinum electrodes, insulated to within 2 mm of the tip, were inserted, in a posterior direction, between whisker rows A/B and C/D of the left whisker pad, ensuring the whole pad received activation following electrical stimulation.

The animal was placed inside a 7 T small bore MRI scanner (Bruker BioSpec Avance II, 70/30, Bruker Biospin GmbH, Ettlingen, Germany) with preinstalled, actively shielded, 200 mm inner diameter, water cooled, gradient coil (Bruker BioSpin MRI GmbH B-GA20. 200 mT/m maximum strength per axis with 200 µs ramps). Inside the magnet bore an electrically filtered and isolated heating blanket (Harvard Apparatus Inc. USA), and rectal probe, maintained body temperature. The animal was artificially ventilated (Zoovent Ltd, UK) with medical grade air and breathing rate measured using a pressure sensitive pad (SAII, USA — Model 1025L Monitoring and Gating System). Standard FLASH based imaging was used to guide subsequent positioning of the fMR imaging plane.

SS-SI-VASO as described above was implemented in Paravision V running Topspin 3.1 (Bruker). Image sequence parameters were TR/TI_1_/TI_2_/TI_3_/TE = 3200/1100/1500/2530/10 ms respectively.A 2 mm thick coronal imaging slice covering the somatosensory whisker barrel cortex; with field of view 30 x 30 mm2 and matrix size 64 x 64 (in-plane resolution 470um2) was used. Use of a coronal imaging plane ensures that the blood refilling conditions (discussed above) are fulfilled and enables depth profiling of response through the cortex.

Intrinsic optical imaging spectroscopy is a well-established technique for delivering high resolution, spatially resolved maps of changes in total haemoglobin (HbT) and oxygen saturation (Chance 1991) (and subsequently deoxy- and oxy-haemoglobin, Hbr and HbO_2_ respectively). A switching galvanometer system (Lambda DG-4 Sutter Instruments Company) with 4 wavelength filters (λ — 495 ± 31, 587 ± 9, 559 ± 16 and 575 ± 14 nm) was used to illuminate the cortex through the medical endoscope. A CCD camera running at 32 Hz (8 Hz effective frame rate for each wavelength) captured remitted/reflected light images with an in-plane spatial resolution of 80µm * 80µm. Light attenuation in response to a stimulation event was calculated and subsequent spectral analysis utilised a modified Beer–Lambert law approach. Mean differential pathlength was estimated, via Monte Carlo simulation of light transport through a parameterised heterogeneous tissue model, assuming a baseline blood volume of 106 µM and mean blood oxygen saturation of 50% (Kennerley et al. 2009). Resultant 2D maps of concentration changes in HbO_2_, Hbr and HbT, alongside raw optical images of the cortical surface aid positioning and cross-validation of image geometry to MRI based on large veins, clearly visible in both modalities (Fig. 1C).

Concurrent OIS and VASO based fMRI data was recorded in response to 16 s electrical stimulation of the whisker pad (5 Hz, 1.2 mA). In all experiments an initial baseline of 60 s was collected, followed by fifteen stimulation events, each with an inter-stimulus interval (ISI) of 86 s. Additional functional experiments used hypercapnic respiratory challenge. Trials were 9 min in duration, consisting of 2 min baseline (medical air), 5% increased FiCO_2_ for 5 min, and a further 2 min baseline. Trials were repeated 12 times to increase CNR.

Following completion of the VASO fMRI experiments, 10 mg/kg of a monocrystalline iron oxide nano-compound (MION) contrast agent (AMI-227 Sinerem; Guerbet Laboratories) was infused intravenously. Infusion took place over 1 hr in steps of 0.1–0.8 ml, a total equivalent to 160 µM concentration and allowed quantification (via high resolution T2 weighted imaging - 256 × 256 pixels, FOV = 30 mm, slice thickness = 1 mm, TR/TE = 1000/15 ms, flip angle = 90°, 2 averages) of baseline blood volume fraction following (Troprès et al. 2001).

All electrical whisker stimulation and hypercapnic challenge studies were repeated after full injection of the contrast agent, to measure MR based changes in CBV following Mandeville et.al. 1998. Functional data were acquired using single shot GE-EPI (raw data matrix = 64 * 64, FOV = 30 mm, slice thickness = 2 mm, TR/TE = 1000/12 ms, flip angle 90°, 10 dummy scans). Here concurrent optical measures are a constant feature between pre/post MION experiments and therefore can be used to help normalise MION and VASO based measures of changes in CBV.

### 2.2. Experimental setup for data acquisition in humans

Data for human experiments were acquired cross-centre (see below), on MAGNETOM 7 T scanners (Siemens Healthcare, Erlangen, Germany) using the vendor-provided IDEA environment (VB17A-UHF). For RF transmission and reception, identical single-channel-transmit/ 32-channel receive head coils (Nova Medical, Wilmington, MA, USA) were used. All scanners used were equipped with a SC72 body gradient coil. Informed written consent was given by all participants.

Human hypercapnia data (n = 5) were acquired at the Max Planck Institute for Human Cognitive and Brain Sciences in Leipzig, Germany. These procedures for human volunteer scanning with respiration challenges were approved by the ethics committee of the University of Leipzig. Respiratory challenges to induce hypercapnia in humans were matched to the design implemented in the rodent experiments e.g. 2 min breathing air, 5 min breathing 5% increased FiCO_2_, and 2 min breathing air again.

Somatosensory stimulation data (n = 9) using abrasive cushions were acquired at SFIM at NIH, Bethesda USA under an NIH Combined Neuroscience Institutional Review Board-approved protocol (93-M-0170) in accordance with the Belmont Report and US Federal Regulations that protect human subjects (ClinicalTrials.gov identifier: NCT00001360). Tapping-induced somatosensory activation followed the same task timing as the rodent experiments (16 s activation with 86 s ISI) but with only 12 repeats (total run duration 18 min 12 s). The task consisted of pinch-like finger tapping and passive touching with an abrasive cushion. The participants were instructed about the task timing inside the scanner bore via a projector and mirror system.

Somatosensory stimulation data (n = 3) using brushes and piezo-electric vibration were acquired at Scannexus (Maastricht, The Netherlands), the corresponding procedures having been approved by the Ethics Review Committee for Psychology and Neuroscience (ERCPN) at Maastricht University, following the principles expressed in the Declaration of Helsinki. Three different stimulation tasks were deployed i) finger tapping, ii) passive finger brushing, and iii) passive somat-sensory stimulation delivered by a piezoelectric vibro-tactile stimulation device (mini PTS system, Dancer Design, UK). All tasks involved the left index finger and thumb. For the passive stimulation, a ceramic mini-PTS stimulator module was held by the participant between the left index finger and thumb. The stimulator module houses a vertically moving aluminium disk (diameter: 6 mm) centered in a static aperture (diameter: 8 mm), delivering a 25Hz vibrotactile stimulus (30s with random ‘silent’ intervals to prevent habituation). The maximum mechanical power delivered to the skin was 75mW (corresponding to a disk movement within the range of 0.5 mm). All stimulation scripts are available for PsychoPy v2.0 and for Presentation v16.1 on Github (https://github.com/layerfMRI/Phychopy_git/tree/master/TappingWithTrTiming and https://github.com/layerfMRI/Phychopy_git/tree/master/Piezo/S1ANFUNCO). Experimental parameters for VASO data acquisition across the experimental paradigms above followed (Huber et al. 2014b) and were in-line with the rodent experiments. In short, to account for short arterial arrival time, the inversion efficiency was reduced to 75%. Further sequence parameters were TR/TI_1_/TI_2_/TE = 3000/765/2265/19 ms. For the hypercapnia data nominal resolution was 1.5 x 1.5 x 1.5 mm3. For laminar comparisons of CBV change in S1, in response to somatosensory stimulation, higher-resolution experiments (Huber et al. 2020b; 2017) were performed with nominal in plane resolution of 0.75 x 0.75 mm2 and slice thickness of 0.7 - 1.8 mm.

### 2.3. General VASO-based data analysis

All fMRI time series were analysed with standard layer-fMRI VASO procedures, as explained in multiple previous publications (Huber et al. 2020a;b; 2017; 2014b; 2018) and our accessible open source online tutorials (https://layerfmri.com/analysispipeline/). In short, time-resolved image repetitions were motion corrected using SPM12 (UCL, UK) (Penny et al. 2007). Volume realignment and interpolation were performed with a 4th-order spline. In order to minimize effects of variable distortions (non-rigid motion), the motion was estimated in a brain mask generated in AFNI (Cox 1996).

Raw VASO data consisted of interleaved acquisitions of MR signal with and without blood nulling (corresponding to CBV and BOLD weighted images). To eliminate unwanted contamination of extravascular BOLD signal within the extracted VASO contrast images the analysis pipeline treated odd and even time points separately for the motion correction and subsequently divided the resultant images by each other (Huber et al. 2014b).

For both rodent and human data, the ensuing layer-dependent signal sampling required manual estimation (based upon T_1_- weighted EPI data) of boundary lines between the grey matter (GM) ribbon, cerebrospinal fluid (CSF) and white matter (WM) to account for cortical curvature. A coordinate system across cortical layers^1^ and columnar structures were estimated in LAYNII (https://github.com/layerfMRI/LAYNII). LAYNII is an open source C++ software suite for computing layer functions (Huber et al. 2020c). We estimated the depth of equi-volume distributed layers (Waehnert et al. 2014). With the resolution of 0.47-0.75 mm, we obtained 4-8 independent data points across the thickness of the cortex. Across these data points, we created 11 layers^2^ across the thickness of the cortex (2mm for both human and rat somatosensory cortex) on a 4-fold finer grid than the effective and nominal resolution.

## 3. Results

### 3.1. Concomitant OIS and VASO

VASO-derived CBV changes and BOLD fMRI maps are shown in Fig. 2, alongside optical measures of the underlying haemodynamics, in response to somatosensory stimulation of the whisker pad. Extracted ROI based time series are shown in Fig. 3. Quantitative estimates of changes in oxygenated haemoglobin(ΔHbO) and total blood volume (ΔHbT) can be made based on the parameterized baseline values (see methods). The sensitivity (in terms of CNR) of all modalities was high enough following trial averaging to detect activation changes in the primary somatosensory cortex. The large draining veins (visible in both structural MRI and raw/analysed OIS data) were used as fiducial markers to align the axial fMRI slice within the field of view of the optical endoscope (turquoise outlines in Fig. 2). Alignment confirmed that in two animals the fMRI slice position overlapped with large pial veins draining the primary somatosensory cortex of the rat brain (top two rows of Fig 2). A BOLD signal change extending away from the somatosensory cortex and towards the superior sagittal sinus is highlighted (black arrow). Interestingly, in one animal, this venous signal (4.1% at yellow arrow in Fig. 2) was greater in magnitude than the signal in the neurally active region within the somatosensory cortex (2.2% at orange arrow in Fig. 2). No corresponding venous signals extending away from the somatosensory cortex were observed in the CBV-based functional maps (even when different thresholds were set). It is noted that due to the position of the rodent head with respect to the static magnetic field the signal in the draining vein (at 90 degrees to the field) is most likely amplified due to the BOLD signal magnetic field angle dependence (Chu et al. 1990)

**Figure 2.**
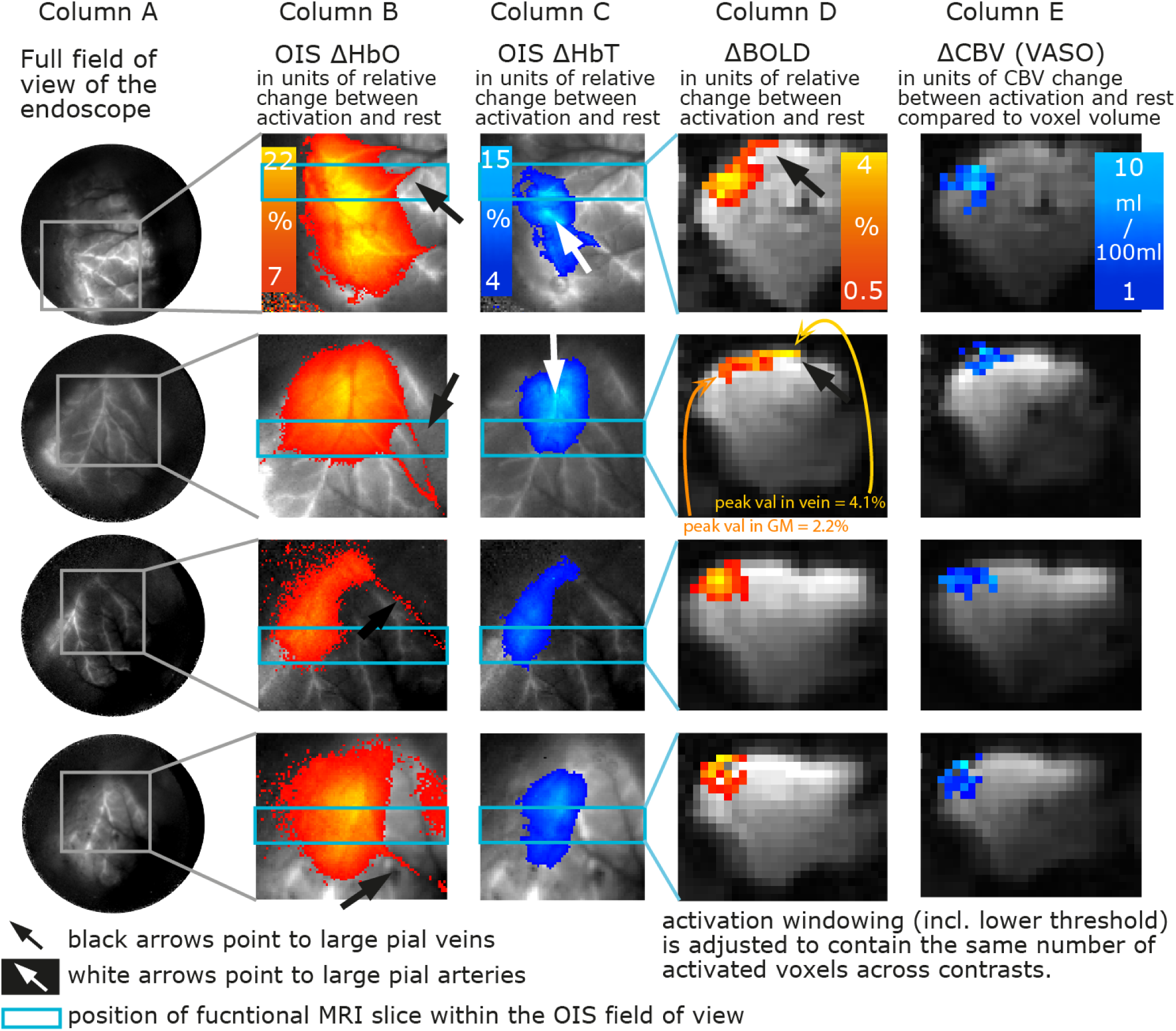
OIS and fMRI activation maps of four animals. Data here shows that VASO-CBV contrast is not sensitive to the large draining veins, which are clearly visible in OIS ΔHbO_2_ and fMRI ΔBOLD activation maps. The VASO CBV response is observed to be specific to the whisker barrel cortex, similar to OIS measures of ΔHbT. Each row depicts one individual animal. **Column A)** depicts the total FOV of the non-magnetic endoscope used to enable concurrent imaging. The grey scale here references HbT response to C_*O*_ 2-induced hypercapnia. Note, arterial vessels are bright, brain tissue is grey, and pial veins display dark contrast. This contrast is used as an underlay/reference image for the somatosensory stimulation results shown. **Columns B)-C)** depict OIS estimates of ΔHbO_2_ and ΔHbT. Draining veins show strong changes in ΔHbO_2_, distant from activated tissue (black arrows). HbT changes, however, are confined to activated tissue. Pial arteries are clearly a major contributor to the HbT change (white arrows). **Columns D)-E)** depict coronal BOLD and CBV fMRI slices extracted from VASO. In animals where the fMRI slice covered the large pial veins (top rows), significant BOLD signal change distal of the barrel cortex (and highest CBV change) was observed (black arrows). Such signals are attributed to venous BOLD weighting amplified by the vessel orientation to the main static magnetic field.

**Figure 3.**
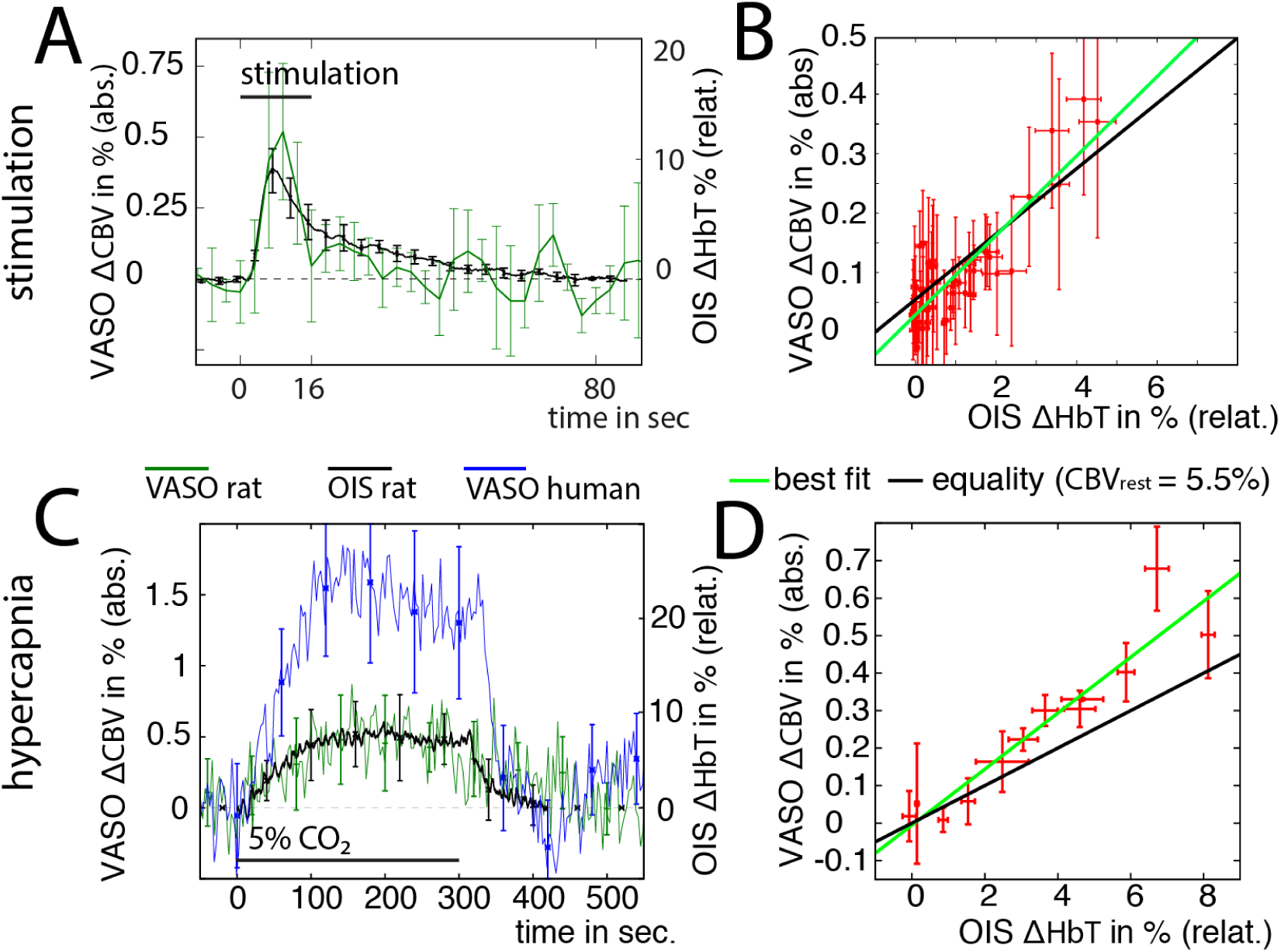
Comparison of OIS and VASO-CBV measurements in time. Unique temporal characteristics of CBV and HbT responses to neuronal activation and respiratory challenges are explored. Panels A) and C) depict CBV and HbT time courses during somatosensory stimulation and hypercapnia, respectively. Human data time series for hypercapnia is shown for completeness (see section 3.3). It can be seen that the time course dynamics between both modalities (VASO and OIS) are in agreement (within standard deviation error). Note that these time-courses show the unique feature of a slow return to baseline after neuronal activation (across a time scale of 80 s) without the post-stimulus undershoot typically observed in equivalent human data. Error bars reflect inter-subject standard deviation across participants. Panels B) and D) show the same data in the form of scatter plots. For the sake of clarity, individual points refer to the averaged signal of multiple adjacent time points. No hysteresis is observed. Data in panels A) and B) refer to ROIs in S1 only, while data in C) and D) refer to the total GM of the right hemisphere within one acquired slice. While both imaging modalities, VASO and OIS, respectively, provide estimates of change in units of %, the reference volume is not the same. Percent volume change in OIS refers to relative (relat.) stimulus-evoked volume changes compared to the volume estimates at baselines (without task). Percent volume change in VASO refers to absolute (abs.) stimulus-evoked volume changes compared to the volume of the imaging voxel (independent of the baseline CBV).

In contrast, in the two animals depicted in the bottom rows of Fig 2, the fMRI slice position does not overlap with large pial veins. In such cases we observed that the activation ‘extent’ of both BOLD and CBV (based on VASO contrast) signal changes were spatially comparable.

In all cases, the optical estimates of total/oxygenated haemoglobin do not show enhanced sensitivity to large draining veins. Indeed HbT changes appear greater in pial arteries (white arrows). These macro-vascular arterial HbT responses are expected also to contribute to the overall layer-dependent activation profiles measured here. Thus, the final VASO and MION layer profiles are expected to reflect both macrovascular and microvascular CBV changes.

Time courses of induced changes in CBV (VASO) and HbT (optical) are shown in Fig. 3A, based on regions of interest drawn from the activation maps in response to somatosensory stimulation shown in Fig. 2. As shown, there is excellent temporal agreement between the two measures. The reader is reminded that VASO CBV is a reflection of the plasma volume, while optical imaging reflects hematocrit volume. Furthermore OIS estimates HbT changes in relative percent changes (to an assumed 106 µM baseline (Kennerley et al. 2005)), while VASO estimates CBV changes in relative percent volume change compared to the voxel volume. To com-pare VASO and OIS reponses in the same *relative* units (of baseline blood volume), the VASO signal was normalized to a literature value baseline CBV, which is assumed to me 5%in GM. This value has been established across two decades of VASO history (Donahue et al. 2006; Gu et al. 2006; Hua et al. 2009; Huber et al. 2014b; Lu et al. 2003; 2013; 2004b;c; Poser and Norris 2007; Scouten and Constable 2008; Shen et al. 2009). The somatosensory CBV changes in response to whisker stimulation peak in the range of 5%-10% (compared to the baseline CBV). Such amplitudes are consistent with previously reported ΔCBV estimates in the pre-clinical domain (Hillman et al. 2007; Kennerley et al. 2005; 2012a; Lee et al. 2001; Tian et al. 2010), whereas they are about 5-10 times smaller than previously reported VASO signal changes in humans (Beckett et al. 2020; Chai et al. 2019; Finn et al. 2019; Guidi et al. 2017; Huber et al. 2018; Kurban et al. 2020; Persichetti et al. 2020; Yang and Yu 2019; Yu et al. 2019).

A scatter plot of group-averaged time points of HbT and CBV change measured via OIS and VASO respectively is shown in Fig 3. The concurrent measures are within standard error of a 1:1 relationship across the whole time series (no hysteresis was observed). Cross modality comparisons of stimulus induced somatosensory activation are somewhat limited by the low sensitivity of VASO and correspondingly large error bars. It is noted that hypercapnia-induced CBV/HbT responses (which were higher in magnitude and longer lasting) yielded more reliable and consistent responses across animals (Fig. 3C-D). The time courses of OIS and VASO were very consistent and both exhibited a characteristic delayed compliance on return to baseline after trial cessation (in-line with Windkessel model predictions (Kida et al. 2007; Kong et al. 2004; Mandeville et al. 1999)).

### 3.2. VASO vs. MION derived CBV and associated cortical profiling

Layer-dependent responses of CBV change estimated with the SS-SI-VASO sequence were compared to layer-dependent responses of MION fMRI in the primary somatosensory cortex. Fig. 4 depicts the respective maps (panel A) of depth-dependent (f)MRI signals and the corresponding group-averaged functional layer-profiles (panels B-C). While GE-BOLD is dominated by the draining vein effect and shows the strongest functional signal changes at the cortical surface, VASO and MION based CBV responses are strongest inside the GM of the activated brain region. Neither CBV measure displayed confounding venous weightings. When analysed in their native physical units, VASO and MION CBV profiles show diverging layer-profiles at the brain surface (pink arrow). The MION CBV profile is dominated by thalamic input layers IV, consistent with previously reported MION-CBV layer profiles in preclinical studies (Goense et al. 2007; Kennerley et al. 2005; Kim and Ugurbil 2003; Mandeville et al. 2001; Silva and Koretsky 2002; Silva et al. 2007; Zhao et al. 2006). The VASO CBV profile, however, also exhibits a similarly elevated CBV response in the superficial layers. This is consistent with previous layer-fMRI VASO studies (Beckett et al. 2020; Chai et al. 2019; Donahue et al. 2017; Finn et al. 2019; Guidi et al. 2017; Huber et al. 2014a; 2015; 2017; 2016; 2018; Kurban et al. 2020; Persichetti et al. 2020; Yang and Yu 2019; Yu et al. 2019) and might appear in conflict with the large body of preclinical studies. Note that the superficial activity in VASO profiles is located within the cortex and is approx 0.8 mm deeper than the GE-BOLD peak response above the cortical surface. While the seemingly different CBV response of VASO and MION fMRI in the superficial layers was an unexpected finding in this study, it can be fully explained by the fact that VASO and MION fMRI estimate CBV changes in different units of percent. While MION estimates CBV changes in relative percent changes compared to the CBV at rest, VASO estimates CBV changes in relative percent volume change compared to the voxel volume. When analyzed in the same physical units (ml blood volume change per 100 ml of tissue) the CBV response in the superficial layers becomes indistinguishable between VASO and MION fMRI. Panel C in Fig. 4 depicts the MION and VASO layer profiles in the same physical units. In order to convert the MION CBV profile from relative percent CBV changes to absolute percent CBV changes, we used the quantitative CBV map of baseline CBV (depicted in panel A) as estimated based on (Troprès et al. 2001)) and undid the normalization of the MION profile, layer-by-layer.

**Figure 4.**
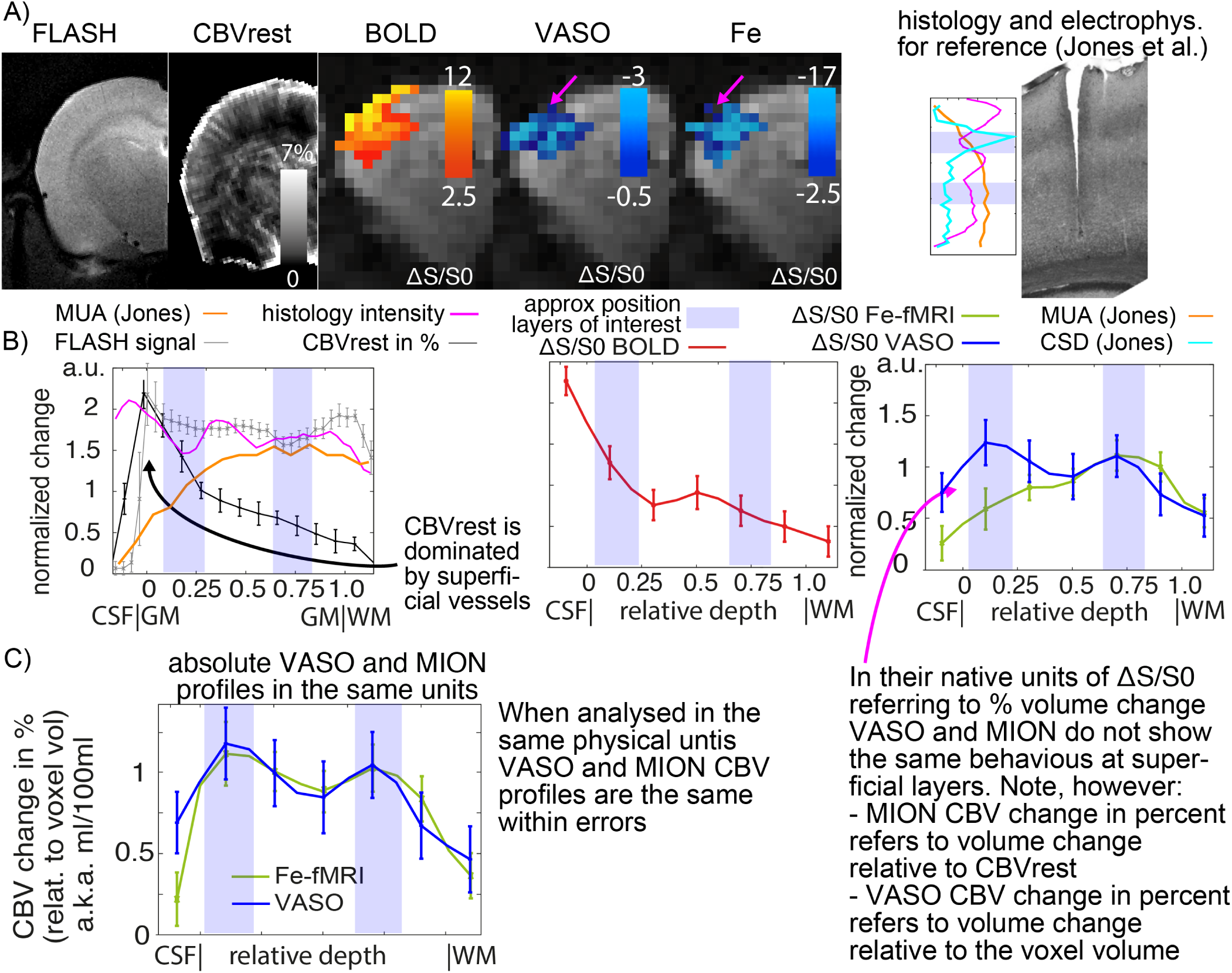
Comparison of CBV layer profiles in the rodent primary sensory cortex with VASO and MION. Data shows that CBV changes estimated from functional VASO and MION maps show the same cortical layer profiles, when analysed in the same physical units. Panel A) depicts the coronal slices from the various MRI contrasts acquired through this study (from one representative animal). High-resolution FLASH data are used to estimate the approximate position of cortical layers (excitatory/inhibitory) (Narayanan et al. 2017). See also included electrophysiology and histology (cytochrome oxidase staining) data from Jones et al. 2004 for the position of the expected excitatory and inhibitory layers. The CSD spike (in rat) is quoted to relate to “primary sensory input from the thalamus” (first bump in the VASO profiles). The MUA peak deeper in the cortex is “likely to reflect the spiking activity of projection neurons in the deeper layers” (second bump in the VASO profiles). The high-resolution CBVrest map is used for retrospective normalizing of CBV change in units of relative changes. The CBVrest was estimated by means of the (Troprès et al. 2001) Vessel size Index model and variable dosage multi-echo FLASH data. Functional activation maps refer to percent signal change of the respective contrast. The activation maps of BOLD signal change are dominated by voxels at the cortical surface (0 mm). Note that the superficial CBV change seems to be stronger in VASO activation maps compared to MION (pink arrow). Panel B) depicts the corresponding layer profiles of all the depicted MRI modalities in the form of profile plots (cortical depth in mm and labelled gray matter, GM, and white matter, WM, where appropriate). The profiles refer to manually drawn ROIs of the primary somatosensory cortex. Signals are averaged within layers and across all animals. The ROI is intended to cover a bit more than GM, only. The closest CSF and WM are also included to see how the signal goes to baseline without partial volumning. Standard errors across subjects are shown. The baseline CBVrest is strongly dominated from the superficial layers (black arrow) reflecting known vascular size/density. As indicated in panel A), VASO and MION profile in their native units of % volume change, diverge at the superficial cortical layers (pink arrow). Panel C) shows that when the same data are evaluated in the same physical units of absolute ΔCBV (ml / 100 ml of tissue), an additional signal component in the superficial layers becomes apparent in the MION data. Only when VASO and MION based fMRI signals are analysed with the same physical units, the resulting layer-fMRI profiles are comparable.

### 3.3. Cross species comparison

Since VASO is a non-invasive contrast, and the resulting CBV measures show improved spatial localisation, with proven little venous weighting compared to BOLD fMRI, it is ideally suited for translational neuroimaging. The VASO ap-proach used here, with the same sequence parameters across species (rat and human), opens up new potential translation of research findings from the preclinical settings to human neuroscience (cognitive research based) for laminar based brain studies. Unfortunately, animal and human research on functional brain mapping to this day remains in separate silos with limited cross-over. Interpretation of human fMR imaging data in terms of neuronal inputs and layer projections would benefit from animal studies in which VASO and electrophysiology could be completed concurrently (Logo- thetis et al. 2001). While VASO may help converge the fields of study, we do of course recognise that there are obvious anaesthetic confounds here, but this is at least a step in the right direction.

Fig. 5 depicts cortical profiles of VASO and BOLD fMRI responses for stimulation in the primary somatosensory cortex (whiskers for rats and fingers for humans) (Arkley 2014; Arkley et al. 2014). The qualitative shape of the profiles are very similar. In both species, BOLD shows the strongest response toward the cortical surface, while VASO shows strongest activation changes within GM. Note that the nominal resolution of the rat results (0.47 mm) is almost double the resolution of the human results (0.75 mm). While the profiles are comparable across species, their respective amplitudes are significantly different. The CBV changes in rats are about 4 times weaker than the CBV changes in humans, which might be related to i) interspecies differences, ii) anesthesia-related differences in physiology, or iii) the specifics of the simulation task. In this study, the experimental setup was chosen to be minimally affected by inter-species and anesthesia effects. Namely, we chose a somatosensory stimulation task over a visual task (used more widely in human neuroscience) to eliminate potential processing pathway bias. In addition we used an anesthesia protocol that has been previously validated with awake animal imaging (Martin et al. 2013).

**Figure 5.**
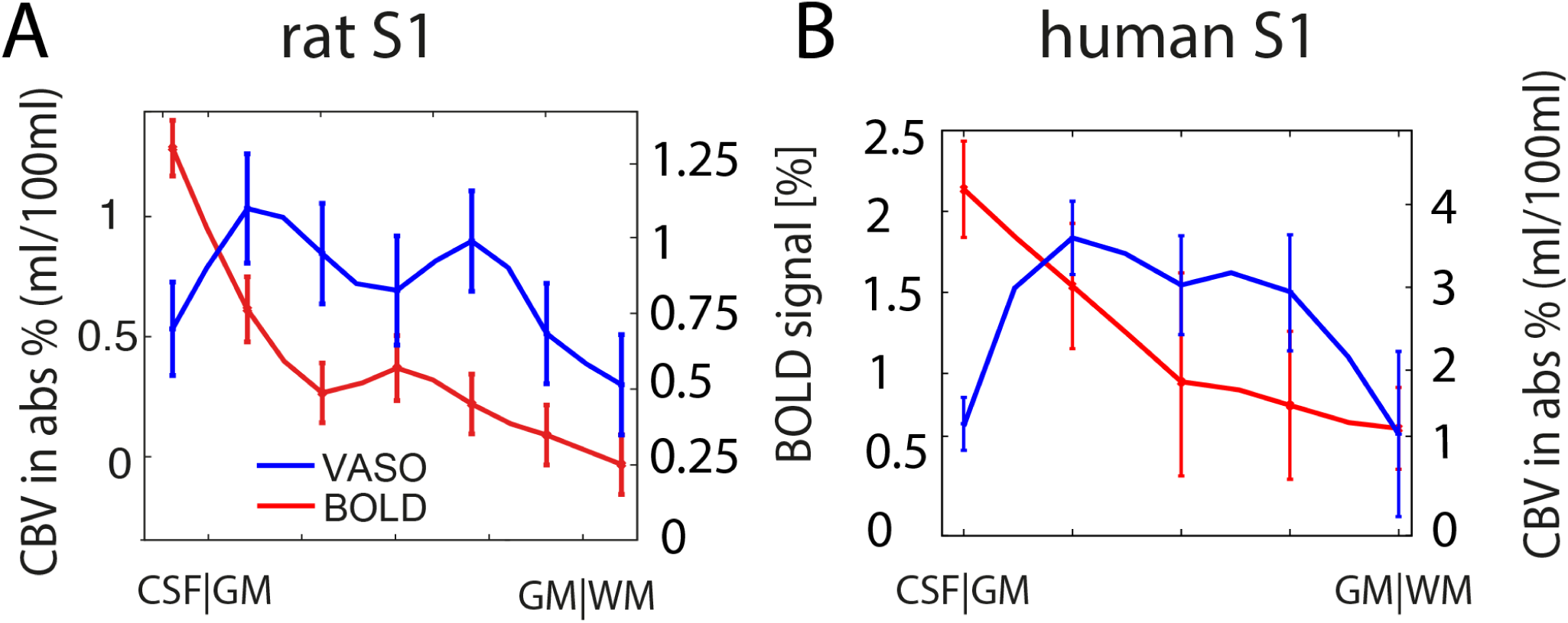
Cortical profiles of BOLD and VASO signal change in rat S1 (A) and human S1 (B). Data shown refer to averages across five rats and nine human participants, respectively. Laminar-dependent profiles reveal different shapes for VASO and BOLD fMRI. BOLD signal change is highly dominated from surface laminae (resulting from known vascular weighting), while VASO peaks in middle cortical laminae, presumably reflecting thalamocortical input, as expected. Note the different scaling of the y-axis in panel A) and B). Error bars refer to standard error of mean across participants.

The somatosensory stimulation task across panels A and B, however, was different. While rats were passively stimulated (sensory only), humans were asked to perform an active tapping movement. This active moment in humans was chosen to maximize the detectability of activation changes, despite the SNR-starved sub-millimeter acquisition.

In order to investigate the influence of active and passive somatosensory stimulations on the magnitude of CBV changes, the experiments were repeated in humans with additional (sensory only) somatosensory stimulation tasks. We measured the CBV response to brushing and piezo-electric vibrations to achieve this. Fig. 6 depicts the respective CBV activation maps and the corresponding group averages of the layer-depend CBV profiles. It can be seen that passive somatosensory stimulation (brushing and piezo) results in 3-4 times smaller CBV responses compared to active finger tapping (similar results are found for respiratory challenge -see Fig. 3C). The magnitude of passive stimulation-induced CBV change in humans is comparable to the response magnitude in rats (horizontal black line in panel C of Fig. 6).

**Figure 6.**
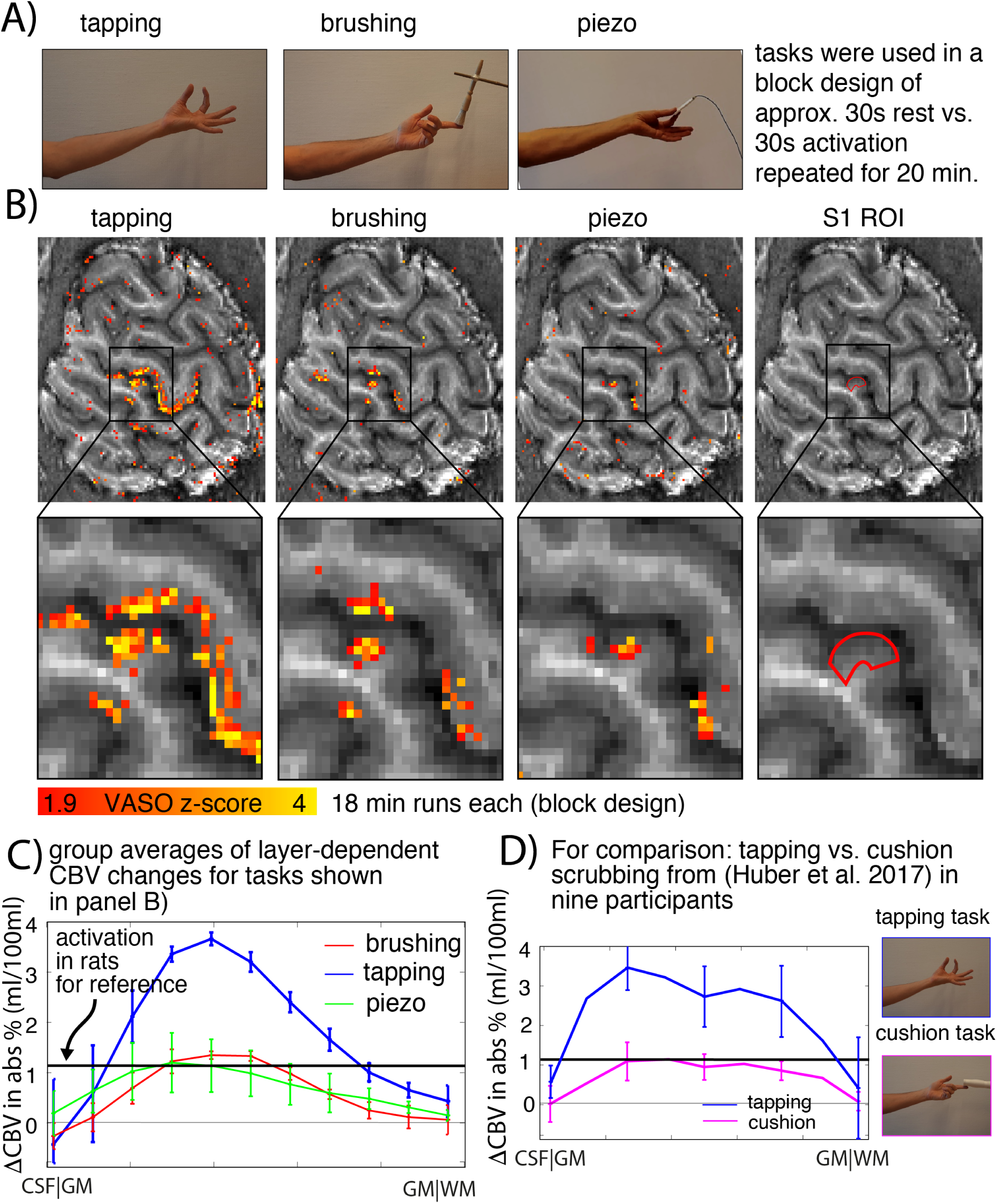
Layer-dependent CBV results for various somato-sensory stimulation paradigms. Data exemplifies that tapping induced activation changes in the primary somatosensory stimulation are significantly stronger than tasks that do not include active movement. This suggests that the different CBV response magnitude between existing rat and human data (often used as evidence to discredit VASO contrast/signal source under-standing) is due to the use of active vs passive sensory tasks. Panel A) depicts the different tasks that are used here to induce activation in the primary somatosensory cortex: active pinch-like finger movement of thumb and index finger, passive brushing of the index finger with a non-metal brush, vibration of the thumb and index finger with a piezo-electric device, and a passive touch with an abrasive cushion (replicated from a previous study (Huber et al. 2019)). Panel B) depicts activation maps of CBV change of a single representative participant. It can be seen that in the index finger region of the primary somatosensory cortex (highlighted in red), the task-induced CBV change is largest for tapping induced activation. Panel C) depicts the corresponding group-averaged layer profiles. While all of the tasks show similar layer-profiles that peak within GM, the overall magnitude is highly variable across tasks. The tapping induced (active) activation is 3-4 times stronger than tasks that are passive sensory trials. Panel D) depicts corresponding results from a previous study (Huber et al. 2019) using a slightly different touching task, with the same conclusion. It is noted that the shape of the profile is highly affected by the effective spatial resolution and corresponding partial voluming effects.

## 4. Discussion

In this study, we compared high-resolution functional brain imaging modalities in the primary somatosensory cortex in both humans and rats. We used these comparisons to validate that the popular VASO contrast is indeed independent of un-wanted signals from large draining veins. This study under-scores VASO’s usability for layer-fMRI application studies in humans.

### 4.1. Validation of CBV weighting in VASO

Previous VASO validation studies in humans have used contrast agent-based methods; including Gd DTPA (Lin et al. 2011), MION (Jin and Kim 2008) and PET-tracers (Uh et al. 2011). CBV-weighting was qualitatively compared by averaging low spatial resolution data across large ROIs covering the whole visual cortex. Following this unrefined approach, VASO signal was found to be qualitatively comparable to CBV changes measured with Gd DTPA across different stimulation frequencies (Lin et al. 2011); but offered little in terms of cortical depth profiling.

In preclinical work, Jin and Kim (Jin and Kim 2008) implemented a version of VASO for ultra high resolution fMRI. Their VASO method provided reliable CBV weighting in deep cortical layers, where CBF effects of macrovasculature and BOLD contaminations are obviously minimized. Here, upon explicit exclusion of signal from inflow-contaminated superficial cortical layers, VASO activation patterns were found to be similar compared to results from iron oxide contrast agent based fMRI signals.

In contrast with the initial VASO studies more than a decade ago, the results shown in the present study are acquired with an adapted VASO method that can account for i) unwanted inflow flow effects in superficial cortical layers and ii) confounding BOLD contamination. Hence, the presented results in this study enable the first quantitative spatio-temporal validation of VASO across cortical depths. Data shown in Figs. 2, 3 and 5 validate the CBV weighting of VASO in reference to the spatial sensitivity of large draining veins and with respect to temporal features such as a slow post-stimulus recurrence to baseline. Furthermore, whole-region VASO-CBV and OIS-HbT measures in response to both neuronal activation and respiratory challenge are of the same magnitude (within error) in terms of their respective quantitative changes (volume fraction vs hematocrit).

### 4.2. Differences between rats and humans

Since VASO is a non-invasive method for CBV mapping, localised to the active brain region and displaying limited macro-vessel weighting, it offers a unique chance for translation between preclinical research and human neuroscience. VASO fMRI allows researchers to compare estimates of CBV cortical layers between humans and animal models by means of the same easily implemented (open source) imaging modality. As such, often used arguments citing discrepancies between human and preclinical results can no longer be attributed to the use of different imaging modalities (e.g. MION vs VASO). Instead, any remaining discrepancies between human and preclinical results in humans and preclinical models can instead be attributed to the important real differences in vascular or neuronal physiology and function. For example, the results shown in Figs. 3C and 5 suggest significantly stronger CBV changes in the primary somatosensory cortex in humans as compared to rats. Such differences are well documented in the literature (Kim and Ogawa 2012). Due to lack of further comparative inter-species data, it was previously suggested that such discrepancies are a result of different imaging modalities, rather than inter-species differences or anesthesia effects (van Zijl et al. 2012); and in some cases such differential findings were used to question the actual signal source of the SS-SI-VASO approach. Here we compared the functional signal magnitude across a wide spectrum of somatosensory stimulation tasks in humans. Common touch tasks are active processes instigated by the participant themselves and therefore, it can be hypothesized, triggering higher-order functions (reading instructions, paying attention to the screen, eliciting a motor task to generate sensory sensation - finger tapping). Somatosensory stimulation in anesthetized rats is entirely passive. In this in-depth study we explored a similarly passive touch stimulus in human participants, providing evidence that different activation strengths are largely due to the specific somatosensory task (either active tapping or passive touch). While finger movement-induced CBV changes in the primary sensory cortex resulted in 3-4 times larger VASO responses compared to rats, a similar passive sensory-only task gave VASO response magnitudes that were identical (within error) across species. Given the similar neurophysiological architecture of the primary somatosensory cortex across species, we believe that interspecies differences in the neural processing of such a passive task can be neglected. In this study, we compare human results obtained in the hand area of the primary somatosensory cortex with rat results in the whisker barrel cortex. While it can be argued that we are therefore investigating different touch sensors and the data is incomparable, we emphasize that the areas chosen, hand/fingers & whisker, are responsible for neural processing of the primary form of active touch in both species, respectively. Thus, these brain systems are entirely suitable for interspecies comparisons (Arkley 2014; Arkley et al. 2014). The stimulation design in the rat experiments was optimized to be minimally affected by the anesthesia protocol. Namely, in a preparatory study, we found that the hemodynamic response to a 5 Hz stimulation frequency was almost identical (indistinguishable within error) between awake and anesthetized rats (see Fig. 7 in (Martin et al. 2006)). Thus, we believe that the anesthesia state is not a relevant source of signal magnitude discrepancies across the human and rat experiments conducted in this study.

### 4.3. Hematocrit

The VASO sequence is expected to be sensitive to changes in **total** blood plasma volume. OIS on the other hand detects direct changes in hemoglobin concentration. This means that the two modalities might capture different components of the cerebral ‘blood volume’ change, if the ratio of haemoglobin volume to plasma volume (commonly referred to as hematocrit level) is not constant during activity. Pericytes are believed to constrict capillary diameter to the point where only plasma perfuses their length. During neuronal activity the pericytes can relax and the capillary dilates permitting red blood cell passage. Capillary recruitment is thought to be an important constituent of neurovascular coupling (Bergers and Song 2005). However, a vast body of literature exists suggesting the effect on hematocrit level is negligible during activity and hypercapnia (Bereczki et al. 1993; Herman et al. 2009; Keyeux et al. 1995; Kleinfeld et al. 1998; Kuschinsky 1996; Pawlik et al. 1981; Villringer et al. 1994). Indeed our data adds to this plethora of data; we demonstrate very close correlation of the VASO derived plasma volume changes and OIS derived total hemoglobin changes. This leads us to question whether we have the resolution, or indeed require such resolution, to be sensitive to capillary recruitment as a driver of significant blood volume changes. Indeed scatter plots comparing the magnitude of concurrently measured CBV and HbT responses, suggest that the change in plasma volume slightly exceeds the change in haemoglobin concentration at the peak of respiratory challenge. This either indicates a contradictory decrease in hematocrit or further depth sensitivity dependency (see section 4.5 below).

### 4.4. Physical units of CBV change in VASO-MION comparisons

In contrast to VASO fMRI, the relative signal change recorded with MION is inherently normalized to the local baseline blood volume. This normalization introduces severe inverse macrovascular weighting at the location of large translaminar arterial and venous vessels. Since the macrovascular baseline CBV can vary up to a factor of five across measured laminae at and below the cortical surface (see Fig. 4B or (Goense et al. 2007; Kennerley et al. 2005; Kim et al. 2013)), the intralaminar microvasculature response in upper cortical laminae appears suppressed in the resulting cortical profiles. The large macrovascular contribution at the surface and the fact that MION-based CBV changes are usually reported in percent change tends to underestimate the actual CBV change (ΔCBV) at the surface. Thus, the widely reported feature that CBV-sensitive iron-oxide-based fMRI shows highest activity in deeper cortical layers (Kim et al. 2013) at the site of thalamocortical input layers does not necessarily suggest high local specificity but might in fact be due to inverse macrovascular weighting in the upper cortical laminae. Furthermore, in MION-based fMRI, the high baseline CBV at the cortical surface can result in low MR signal intensity and might thus not capture the relative functional signal change under the Rician noise floor with sufficient statistical detection power.

In this study, for the sake of quantitative comparison of VASO and MION results, we unnormalized the relative CBV signal change by modelling the baseline CBV at rest. This allowed us to compare the respective modalities in the same physical units of ml per 100 ml of tissue. Following such remapping, the cortical layer profiles of VASO and MION matched within error (see Fig. 4C) and did not show the erroneous fMRI signal suppression at the superficial layers. Indeed the characteristic bumps noted in the cortical profiles could correspond to excitatory and inhibitory processing layers (Narayanan et al. 2017) in line with electrophysiology CSD and multi-unit activity respectively (Jones et al. 2004).

### 4.5. Physical units of CBV change in VASO-OIS comparisons

Unlike MION MRI, OIS does not provide estimates about the baseline CBV. Thus, it was not straightforwardly possible to compare VASO with OIS results in VASO’s native units of Δml per 100 ml of tissue. For this reason, using the large body of VASO literature, we converted the raw quantitative VASO signal change into relative ΔCBV changes by means of normalizing it to an assumed value of CBV_*rest*_ (Donahue et al. 2006; Gu et al. 2006; Hua et al. 2009; Huber et al. 2014b; Jin and Kim 2008; Lu et al. 2003; 2013; 2004b;c; Poser and Norris 2007; Scouten and Constable 2007; 2008; Shen et al. 2009; Uh et al. 2011; Wu et al. 2010). Consistently with these earlier VASO studies, a value of CBV_*rest*_ = 5.5% was assumed. Even though the corresponding VASO-Δ *CBV* results are within error to OIS-ΔHbT results, for a slightly larger value of CBV_*rest*_ = 6.5%, the quantitative estimates of VASO-ΔCBV and OIS-ΔHbT results would be identical (see green best fit line in Fig 3). The result that an optimal correspondence between VASO CBV and OIS HbT would be found for slightly larger assumed values of CBV_*rest*_ might be related to the fact that the OIS is mostly sensitive to superficial cortical layers, which have a higher vascular density than deeper layers and are thus above the assumed 5.5% value (see Fig. 4B).

## 5. Conclusion

We conducted a validation study of layer-dependent VASO fMRI by comparing the SS-SI-VASO MR sequence with established preclinical CBV/HbT imaging modalities (contrast-agent based fMRI and intrinsic optical imaging respectively). We found excellent agreement in terms of both spatial and temporal dynamics between all modalities tested in a rat model. Comparing plasma volume measurements and haemoglobin total changes suggested that over the activation period (of somatosensory stimulation or respiratory challenge) there is no change in the hematocrit. In addition we explored the cortical layer dependency of the CBV responses to somatosensory neuronal activity; comparing preclinical rat data with equivalent data from human participants. We found that for quantitative imaging of task-induced CBV changes, it is vital to consider the signal change in the same physical units of ml per 100 ml of tissue. When layer-dependent CBV changes are analyzed in the same physical units, the VASO method provides results that are identical to those obtained from long established preclinical methods based on the infusion of a paramagnetic contrast agent. We found that the resultant CBV response provides better spatial localisation than the equivalent BOLD signal (often weighted by venous macro-vessels), with profile peaks corresponding to excitatory and inhibitory cortical pathways. We also found that the CBV response magnitude depends on whether the stimulus is active or passive. Upon harmonizing the somatosensory stimulation to be a passive sensory-only task in both species, the VASO results were found to be comparable across species. This has important consequences for translation of experimental design between preclinical and clinical neuroscience. Finally, this study provides direct evidence that the non-invasive SS-SI-VASO sequence can map CBV quantitatively, and can provide reliable estimates of CBV change at the spatial scale of cortical layers. Indeed with CBV presenting as a more robust correlate of neuronal activity, maintaining signal change even in light of significant physiological changes (which have been demonstrated to dramatically affect BOLD fMRI (Kennerley et al. 2012a)); VASO fMRI should become the method of choice for high resolution layer dependent fMRI studies in neuroimaging.

## Funding

Laurentius Huber and Aneurin J Kennerley received funding from the York-Maastricht partnership for this project. Laurentius Huber was funded form the NWO VENI project 016.Veni.198.032 for part of the study. Benedikt Poser is partially funded by the NWO VIDI grant 16.Vidi.178.052 and by the National Institute for Health grant (R01MH/111444) (PI David Feinberg). Rainer Goebel is partly funded by the European Research Council Grant ERC-2010-AdG 269853 and Human Brain Project Grant FP7-ICT-2013-FET-F/604102. We thank Harald E Möller and the NMR-group of the MPI-CBS for supporting the part of this study that was conducted at the MPI in Leipzig. The preclinical studies presented here were supported by a United Kingdom Medical Research Council (MRC) grant MR/M013553/1 and Wellcome Trust (WT) Grant 093069/Z/10/Z.

## Animal Husbandry

We thank the technical team within the biological services unit at the University of Sheffield for animal husbandry support.

## Data Acquisition

We thank Kenny Chung for radiographic assistance with experiments conducted at NIH. We thank FMRIF (especially Sean Marrett), and Scannexus (especially Chris Wiggins), where MBIC data were acquired. We thank Dimo Ivanov for the setup of the Hypercapnia delivery system used for the experiments in humans.

## Safety supervision of human experiments

We thank Dr. Ilona Henseler, Dr. Elisabeth Roggenhofer, and Dr. Stephan Kabisch for medical supervision of experiments with hypercapnia. We thank Chris Wiggins, and Bianca Linssen for help with the safe application of human finger brushing inside the scanner bore.

## Financial interest

Rainer Goebel works for Brain Innovation and has financial interest tied to the company.

## Data and Software availability

The data shown here have been acquired over a period of four years across four institutions with variable data sharing policies. All processed human and rat data are available upon request in anonymized form. Data presented in Fig. 5 are already publicly available from previous publications. Data presented in Fig 6 have been acquired under the York-Maastricht partnership and are available as raw data on Openneuro. We are happy to share the used sequence with the vendor approval for Bruker Paravision (versions 3 to 5), as well as SIEMENS VB17B.

## Footnotes

1 Note on terminology: In the context of fMRI, these estimates of cor-tical depth do not refer to cytoarchitectonically defined cortical layers (https://layerfmri.com/terminology).

2 The number of twenty layers was chosen based on previous ex-perience in finding a compromise between data size and smoothness (https://layerfmri.com/how-many-layers-should-i-reconstruct).

## Notes

### Competing Interest Statement

Rainer Goebel works for Brain Innovation and has financial interest tied to the company.

